# TGF-β-R2 is required for HBP induced acute lung injury and vascular leakage for TGF-β/Smad/Rho signaling pathway activation

**DOI:** 10.1101/2022.01.14.476433

**Authors:** Zixuan Liu, Mingming Chen, Yini Sun, Xu Li, Liu Cao, Xiaochun Ma

## Abstract

Heparin-binding protein (HBP), as a granule protein secreted by polymorphonuclear neutrophils (PMNs) participates in the pathophysiological process of sepsis. It has been reported that HBP is a biomarker of sepsis, which is related to the severity of septic shock and organ dysfunction. HBP binds to vascular endothelial cells as one of the primary target sites. However, it is still unclear whether HBP-binding protein receptors exist on the surface of ECs. The effect of HBP on vascular permeability in sepsis and its mechanism needs to be explored. We conducted *in vivo* and *in vitro* study. We demonstrated that HBP bound to transforming growth factor-β receptor type 2 (TGF-β-R2) as a ligand. GST pull-down analysis reveals that HBP mainly interacts with the extracellular domain of TGF-β-R2. HBP induced acute lung injury (ALI) and vascular leakage via activation of TGF-β/SMAD2/3 signaling pathway. Permeability assay suggests TGF-β-R2 is necessary for HBP-induced increased permeability. We also defined the role of HBP and its potential membrane receptor TGF-β-R2 in the blood-gas barrier in the pathogenesis of HBP-related ALI.

## Introduction

Sepsis is an imbalance of systemic inflammation response caused by infection, resulting in fatal organ dysfunction. The release of inflammatory mediators leads to uncontrolled capillary leakage, which is one of the main causes of multiple organ dysfunction syndrome (MODS) and even death in sepsis[1]. At the early stage of inflammation, polymorphonuclear neutrophils (PMNs) are recruited to the site of infection. The activation, chemotaxis, and migration of PMNs lead to vascular endothelial barrier damage, which is closely related to the action of proteins secreted by PMNs. Heparin-binding protein (HBP), also called cationic antimicrobial protein of 37 kDa (CAP37) or azurocidin, is secreted by PMNs as an important granule protein. It is involved in the pathophysiological process of sepsis[2]. HBP is synthesized in advance and stored in secreted particles and azophil particles of PMNs. When the body is invaded by pathogens, HBP is released into the blood to plays an anti-inflammation role. HBP causes vascular leakage by increasing endothelial permeability [3]. Dysregulated HBP level may cause tissue edema, hypotension, and organ dysfunction[4]. In recent years, studies have revealed that HBP is related to the severity of sepsis and can be used as a biomarker[5, 6]. HBP docks on the surface of vascular endothelial cells (ECs) and causes rearrangement of cytoskeleton and vascular leakage through protein kinase C (PKC)[7] and rho kinase signaling pathways[8]. The surface structures of endothelial cells are very complicated. The polysaccharide structure of glycocalyx on the surface of endothelial cells are considered as the HBP-binding site[9]. However, there is no direct evidence that this binding is associated with increased endothelial permeability.

Interestingly, HBP promotes cell migration in many different cells, including fibroblasts, smooth muscle cells and corneal epithelial cells, which plays a role in the process of tissue damage and repair[10–13]. Though the underlying mechanisms remain unclear, this biological behavior is very similar to epithelial-mesenchymal transition (EMT). EMT is an evolutionary conserved biological process that typically takes place during organism development and wound healing process. It causes a switch from epithelial phenotype to the mesenchymal phenotype. Morphologically, cells undergoing EMT are manifested by loss of cell polarity, changes of intercellular junctions, and remodeling of the cytoskeletal architecture leading to the detachment of cells from basement membrane. In addition, EMT also plays an important role in cancer progression.

Transforming growth factor-β (TGF-β) is an evolutionarily conserved polypeptide family, which regulates embryogenesis and tissue homeostasis. TGF-β family members are associated with the pathophysiological mechanisms of many diseases[14]. TGF-β is one of the inducers of EMT process. TGF-β signaling pathway also participates in infectious diseases[15, 16]. Binding of TGF-β to TGF-β receptor 2 (TGF-β-R2), through the canonical pathway, activates transcription factors, which positively or negatively regulate the transcription of target genes, thereby influencing the repair process and EMT[17]. Through the noncanonical pathway, TGF-β-R2 directly regulates the Rho-associated protein kinase (Rho-ROCK) pathway, eventually resulting in actin cytoskeleton reorganization[18]. Activated TGF-β signaling pathway also induces dissociation of adherens junctions (AJs), which promotes tumor metastasis[19].

There are few investigations on whether HBP-binding protein receptors exist on the surface of ECs. The underlying mechanisms of HBP-induced increased endothelial permeability in sepsis remains unclear. Previous study have shown that TGF-β is also a kind of heparin binding proteins[20]. Therefore, we hypothesize that TGF-β-R2 may be a potential receptor of HBP on the surface of ECs. HBP induces acute lung injury (ALI) and vascular leakage upon the binding to TGF-β-R2. Therefore, we conducted the following *in vivo* and *in vitro* studies.

## Results

### HBP bound to TGF-β-R2 as a ligand

The Discovery Studio (DS) 2016 ZDOCK program indicated possible HBP-TGF-β-R2 binding pose in 3D space (Fig. 1A). The H-bond interaction of HBP and TGF-β-R2 were also analyzed using ZDOCK (Fig. 1B). HBP Tyr81, Ser41, Arg23, Asn172, and Ser133 formed H-bonds with TGF-β-R2 (Fig. 1C). These data suggested that HBP bound to TGF-β-R2. Then we confirmed the interaction between HBP and TGF-β-R2 by immunoprecipitation (Fig. 1D). Under a microscope, co-localization of HBP and TGF-β-R2 was observed in HUVECs treated with HBP 10 μg/mL for 30 min (Fig. 1E). We overexpressed TGF-β-R2 in E. coli (BL21), and purified TGF-β-R2 protein to co-precipitate it with recombinant HBP protein. The results showed that there was a direct binding between HBP and TGF-β-R2 (Fig. 1F).

**Fig. 1.**
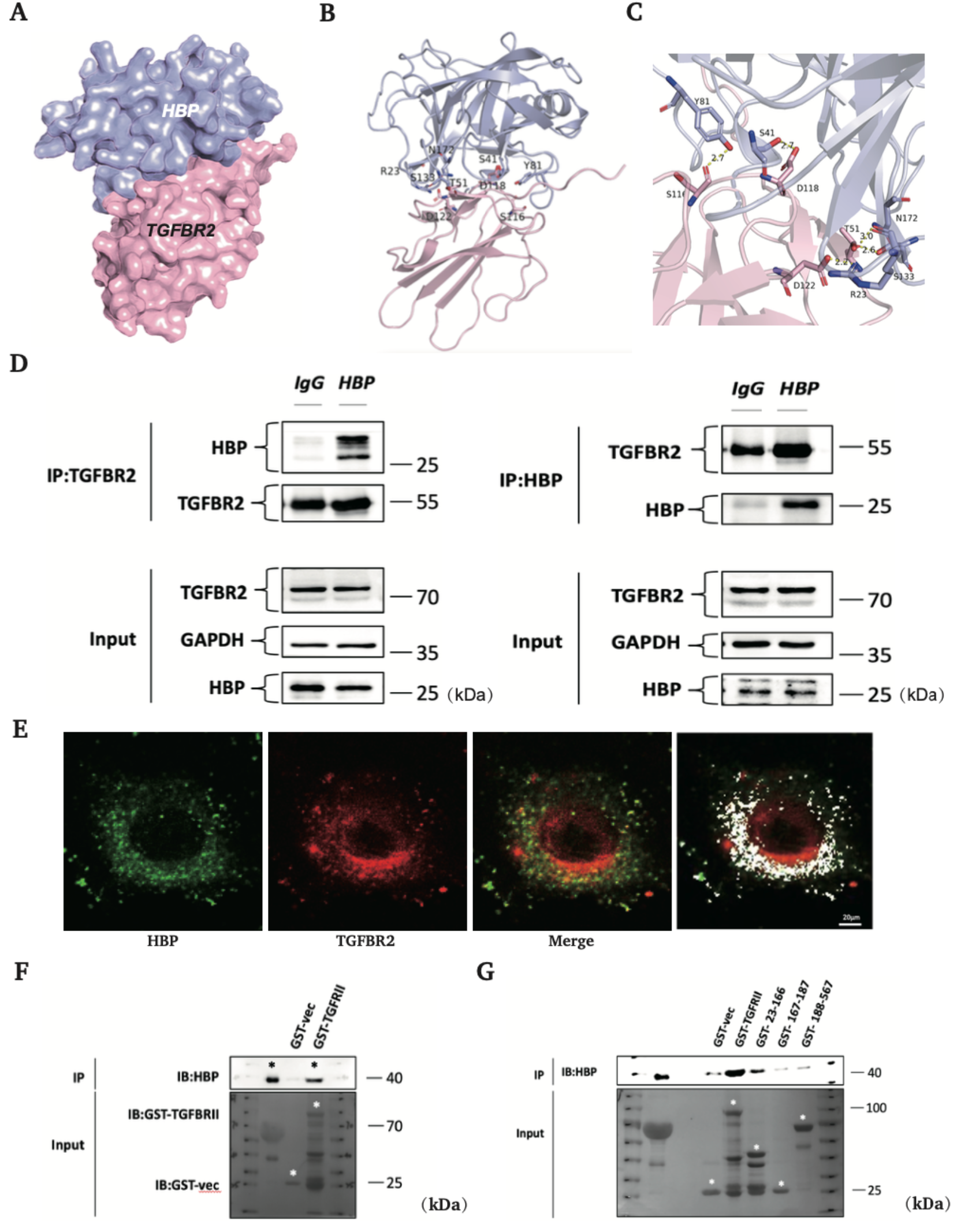
The interaction between HBP and TGF-β-R2. (A) Structure of HBP bound to TGF-β-R2. HBP is shown in blue, and TGFBR is shown in pink. (B, C) HBP-TGF-β-R2 interaction residues. HBP residues are shown in blue, and TGF-β-R2 residues are shown in pink. Hydrogen bonds are shown as yellow dashed lines. The images were constructed using ZDOCK program. (D) Coimmunoprecipitation (co-IP) and Western blot (IP-western) using anti-HBP or anti-TGF-β-R2 antibody or negative control IgG and Protein A/G immunoprecipitation magnetic beads followed by anti-TGF-β-R2 or anti-HBP Western blot to verify endogenous interaction between HBP and TGF-β-R2. (E) HUVECs were grown in medium with 10% FBS for 24 h and were treated with 10 μg/ml HBP for 30 min. the colocalization of HBP and TGF-β-R2 was detected by immunofluorescence using the anti-HBP (primary), anti-TGF-β-R2 (primary), and anti-mouse/rabbit Alexa Fluor (secondary) antibodies. (F, G) Indicated truncates of TGF-β-R2 were constructed according to their functional domains. Transfection of E. coli. (BL21) with the indicated truncates of TGF-β-R2. Coimmunoprecipitation (co-IP) and Western blot (IP-western) using GST Sepharose Beads followed by anti-HBP Western blot to verify the direct interaction between HBP and TGF-β-R2.

We further explored the binding region of HBP and TGF-β-R2. Because TGF-β-R2 is a transmembrane protein receptor, we constructed the truncated plasmid of TGF-β-R2 according to its functional domains, including the extracellular domain of TGF-β-R2, the transmembrane domain of TGF-β-R2, and the intracellular domain of TGF-β-R2. Next, three fragments of TGF-β-R2 were overexpressed in E. coli (BL21). The purified protein fragment was used for co-precipitation with recombinant human HBP protein. The results showed that HBP mainly interacted with the extracellular domain of TGF-β-R2 (Fig. 1G).

### HBP participated in the activation of the TGF-β/SMAD2/3 signaling pathway

The relationship between HBP and the TGF-β pathway has not been reported before. SMAD2/3 is a downstream signaling molecule of TGF-β pathway. Compared with the control group, the protein levels of phospho-SMAD2/3 were significantly upregulated after stimulation with HBP 10 μg/mL for 3 h, 6 h, and 9 h, but there was no change in total SMAD2/3 levels (Fig. 2A).

**Fig. 2.**
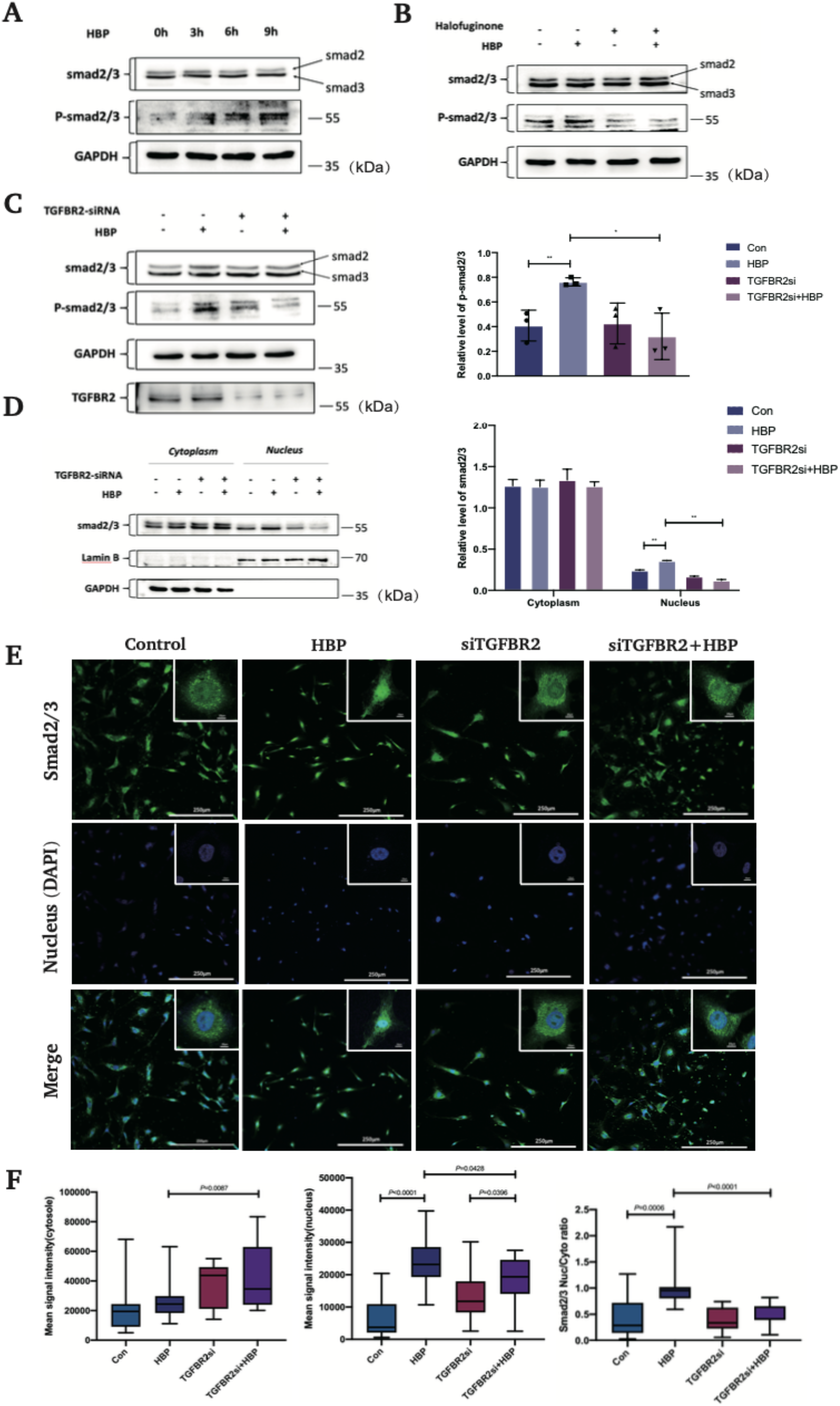
HBP activated TGF-β signaling pathway. (A) HUVECs were treated with HBP at the concentrations of 10 μg/mL for 3 h, 6 h, 9 h and the expression of Smad2/3 and phospho-Smad2/3 was analyzed by Western blot. GAPDH was used as a loading control. (B) Pretreatment of HUVECs with halofuginone (Smad3 inhibitor) at the concentration of 10 ng/mL for 24 h, HUVECs were then treated with HBP for 3-6 h at the concentration of 10 μg/mL and the expression of Smad2/3 and phospho-Smad2/3 was analyzed by Western blot. GAPDH was used as a loading control. (C) TGF-β-R2-siRNA transfected HUVECs were treated with HBP for 3-6 h at the concentration of 10 μg/mL and the expression of Smad2/3 and phospho-Smad2/3 was analyzed by Western blot. GAPDH was used as a loading control. (D) TGF-β-R2-siRNA transfected HUVECs were treated with HBP for 3-6 h at the concentration of 10 μg/mL and were subjected to Cytoplasmic and Nuclear Fractionation assay. The expression of Smad2/3 and phospho-Smad2/3 was analyzed by Western blot. GAPDH was used as a loading control. Protein expression and the nuclear translocation of Smad2/3 in HUVECs by confocal immunofluorescence analysis. (E) TGF-β-R2-siRNA transfected HUVECs were treated with HBP for 3 h at the concentration of 10 μg/mL. HUVECs were then stained against Smad2/3 and DAPI. (F) The nuclear to cytoplasmic ratio of mean Smad2/3 signal intensity. (G) Mean Smad2/3 signal intensity in the cytosol and (H) nucleus. Data were expressed as the mean ± SD (****P < .0001, ***P < .001, **P < .01 and *P < .05, Student’s t-test).

To clarify the key steps involved in HBP-induced activation of the TGF-β pathway, we first divided the cells into four groups: control group, HBP group, TGF-β-R2 siRNA group, and TGF-β-R2 siRNA + HBP group. Compared with the control group, the levels of phospho-SMAD2/3 were increased in the HBP 3h and 6h group, while there were no significant differences in the levels of phospho-SMAD2/3 between TGF-β-R2 siRNA group and TGF-β-R2 siRNA + HBP group. The results indicated that HBP upregulated the phosphorylation of SMAD2/3 through TGF-β-R2 (Fig. 2C). Next, we used halofuginone, a SMAD3 inhibitor[21]. Cells were divided into four groups: control group, HBP group, halofuginone group, and halofuginone + HBP group. Cells were pretreated with 10 ng/mL halofuginone for 24 h before stimulation with HBP 10 μg/mL for 3 h. Western blot analysis was used to detect the expression of SMAD2/3 and phospho-SMAD2/3 protein. As shown in Fig. 2B, there was no significant change in the SMAD2/3 protein level in all groups, while the protein level of phospho-SMAD2/3 in the HBP group was significantly higher than that in the other three groups. Therefore, activation of the TGF-β pathway by HBP was dependent on TGF-β-R2 and SMAD3.

We then attempted to explore whether SMAD2/3 complexes translocated from cytoplasm to nucleus upon stimulation with HBP 10 μg/mL for 3 h. We used western blot assay to verify the levels of SMAD2/3 protein in nucleus and cytoplasm. The results showed that there was no significant difference in the protein levels of SMAD2/3 in cytoplasm among all groups. However, the SMAD2/3 protein levels in nucleus in the HBP group increased significantly, although similar levels were observed in the control, TGF-β-R2 siRNA and TGF-β-R2 siRNA + HBP groups (Fig. 2D). To visualize the effects of TGF-β pathway in HBP-induced SMAD2/3 nuclear translocation, immunofluorescence assay was used to examine nuclear localization of SMAD2/3. As predicted, HBP treatment induced the nuclear accumulation of SMAD2/3, while TGF-β-R2 silencing significantly inhibited this effect (Fig. 2E, F). Thus, we demonstrated that the binding of HBP to TGF-β-R2 is critical for activation of the TGF-β signaling pathway.

### HBP destroyed endothelial barrier through the activation of TGF-β signaling pathway

To further evaluate the effect of HBP-induced-activation of TGF-β signaling on vascular permeability, we conducted the following studies. We divided HUVECs into four groups: control group, HBP group, TGF-β-R2 siRNA group, and TGF-β-R2 siRNA + HBP group. The permeability of HUVECs was detected by trans-endothelial permeability assays. The results showed that the concentration of FITC-dextran (molecular weight ≈ 40kDa) detected in the medium of the lower chamber in the HBP group was higher than that in the other three groups. The results showed that HBP treatment significantly increased the permeability of HUVECs, which was higher than that of TGF-β-R2 siRNA+HBP group. The results suggested that knockdown of TGF-β-R2 could partially antagonize the effect of HBP on endothelial permeability (Fig. 3A).

**Fig. 3.**
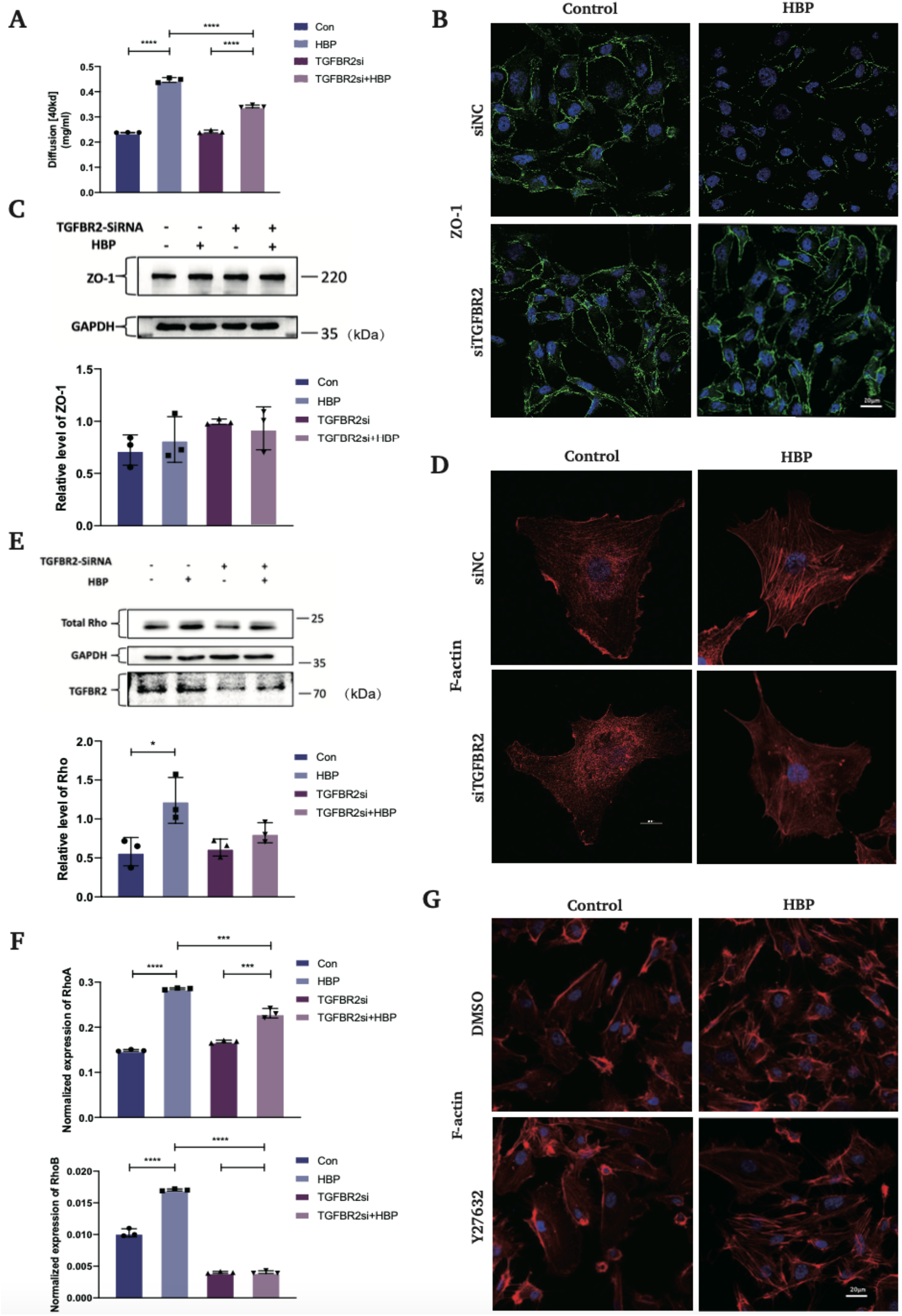
HBP destroyed endothelial barrier through the activation of TGF-β signaling pathway. (A) HBP increases endothelial permeability through TGF-β signaling pathway. TGF-β-R2-siRNA transfected HUVECs were treated with HBP at the concentration of 10 μg/mL for 1.5 h. Permeability of stimulated monolayers is shown as the concentration of FITC-dextran. TGF-β-R2-siRNA transfected HUVECs were treated with HBP for 12 h at the concentration of 10 μg/mL. HUVECs were then subjected to (B) immunofluorescence assay and (C) western blot assay to examine ZO-1. GAPDH was used as a loading control. (D) The same cells as in (B) were subjected to immunofluorescence assay to examine actin cytoskeleton. (E, F) The same cells as in (B) were subjected to RT-PCR and western blot assay to examine Rho. (G) Pretreatment of HUVECs with Y27632 (ROCK inhibitor) at the concentration of 10 μM. The inhibitors were added for the last hour of the 12 h HBP incubation. HUVECs were then treated with HBP 10 μg/mL for 12 h and were subjected to immunofluorescence assay to examine actin cytoskeleton. Data were expressed as the mean ± SD (****P < .0001, ***P < .001, **P < .01 and *P < .05, Student’s t-test).

To further explore the mechanism of HBP on endothelial permeability, we focused on the changes in tight junction (ZO-1) and cytoskeleton. As shown in Fig. 3B, ZO-1 formed a continuous line at the cell–cell junctions, with occasional gaps. After HBP stimulation for 24 h, immunofluorescence assay revealed that the distribution of ZO-1 was fragmented, indicating that tight junction integrity was disrupted. However, ZO-1 protein levels remained unchanged in western blot assay under HBP treatment (Fig. 3C). Then, the actin cytoskeleton organization was assessed by immunofluorescence using rhodamine phalloidin. HBP stimulation resulted in actin reorganization with the formation of thickened and disordered stress fibers in HUVECs. However, there was no significant changes in the control group, TGF-β-R2 siRNA group, and TGF-β-R2 siRNA+ HBP group (Fig. 3D). We also observed that the protein level of Rho was upregulated in HUVECs stimulated with HBP 10 μg/mL for 6 h (Fig. 3E). Next, we attempted to analyze the effects of HBP treatment on the mRNA expression of RhoA and RhoB in HUVECs using qRT-PCR. The results showed that the mRNA levels of RhoA and RhoB were upregulated by HBP (Fig. 3F). TGF-β-R2 silencing and the use of Y27632(ROCK inhibitor) 10 μM could antagonize this effect (Fig. 3G). These results indicated that HBP increased HUVEC permeability by binding to TGF-β-R2. HBP activated Rho-ROCK pathway through TGF-β signaling, resulting in the reorganization of cytoskeleton and the redistribution of tight junction protein (ZO-1).

### HBP induced ALI in mice

Pulmonary edema is one of the indicators of the blood–gas barrier damage, which is usually presented as an increased in the W/D weight ratio of lung tissue. Our study revealed that HBP treatment resulted in significantly higher W/D weight ratios. TGF-β-R2-siRNA treatment markedly reduced the W/D weight ratios (Fig. 4A). Then we investigated the histological changes of lung tissue. HBP administration induced characteristic histological features of ALI, including thickened alveolar septa, vascular congestion, and neutrophil infiltration in the alveolar space. TGF-β-R2 knockdown partially alleviated these pathological changes (Fig. 4B). The lung injury scores were higher in HBP group than that in control group and TGF-β-R2-siRNA+HBP group, indicating that TGF-β-R2 knockdown partially protected HBP-induced ALI. The degree of ALI was further evaluated by EM. SEM showed alveolar protein deposition and alveolar structures destruction. SEM also revealed damage of the blood-gas barrier (Fig. 4C). By TEM, we observed the destruction of membrane contacts, leading to the formation of clefts between membrane structures after HBP stimulation. Disorganization of blood–gas barrier structures was also evident (Fig. 4D). TGF-β-R2-siRNA treatment alone did not affect the structure of the blood–gas barrier. TGF-Β-R2-siRNA significantly alleviated HBP-induced ALI. These above results showed that the injurious effect of HBP was antagonized by TGF-β-R2-siRNA treatment, indicating that HBP-induced ALI via activation of the TGF-β signaling pathway.

**Fig. 4.**
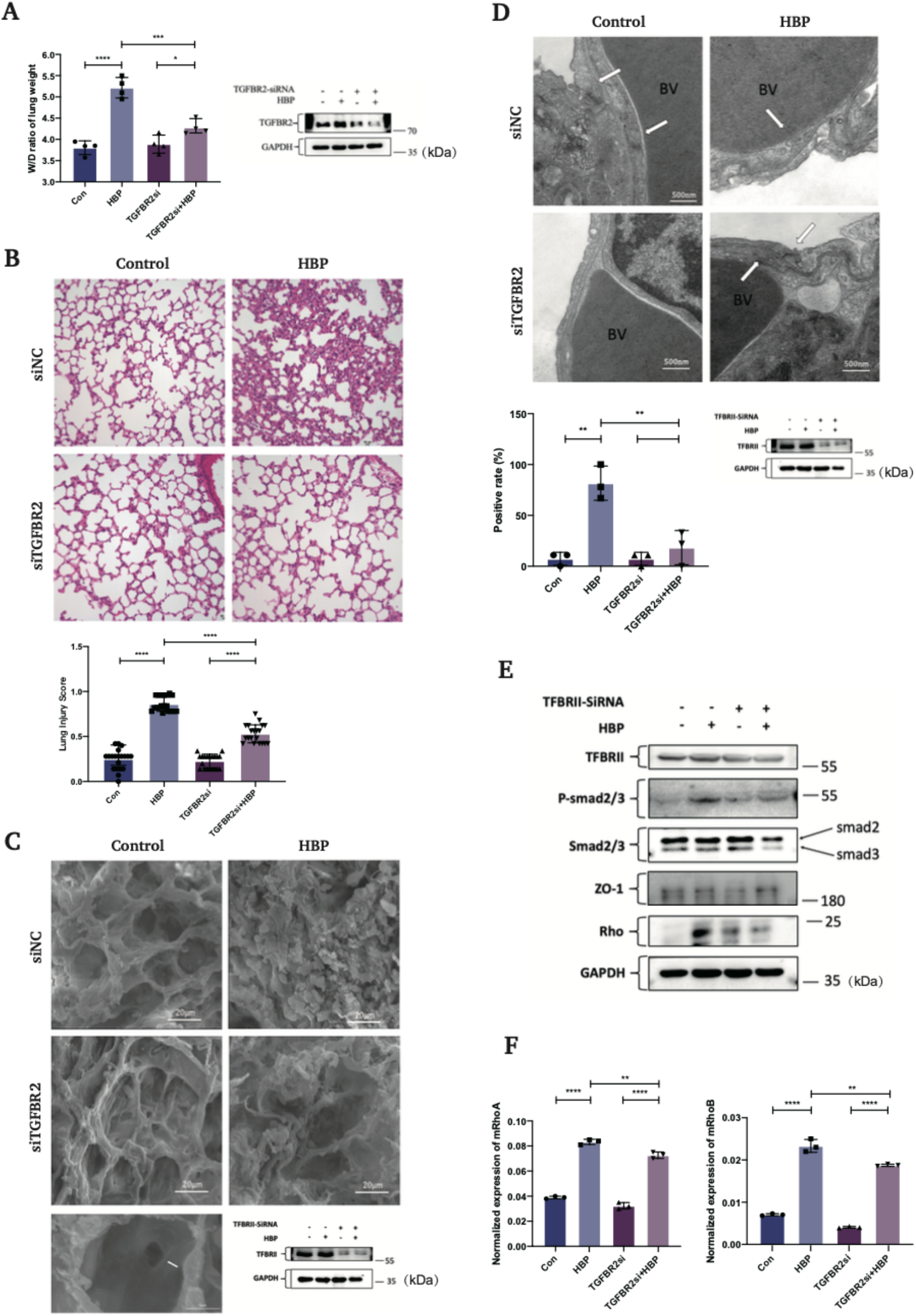
HBP activated TGF-β signaling pathway in the lung tissue in mice. TGF-β-R2-siRNA transfected C57BL/6 mice were treated with HBP intravenously at the concentration of 150 μg/mouse for 6 h. After the treatments, the lung tissue was taken for western blot, RT-PCR. Then the lung tissue was used for wet-to-dry ratio of lung weight analysis, H&E staining, scanning electron microscopy (SEM), and transmission electron microscopy (TEM). (A) Effects of HBP treatment on pulmonary edema were determined by the wet-to-dry (W/D) ratio of lung weight (n=4 in each group). (B) The lungs were stained with hematoxylin and eosin (original magnification × 200), analyzed by SEM (C), arrow shows the destroy of the blood-gas barrier. (D) The lungs were analyzed by TEM. Positive rate referred to the ratio of opened cell junction number over total field vision number. Lung injury scores were evaluated using the lung injury scoring system described in Materials and Methods. (E) The expression of Smad2/3, phospho-Smad2/3, ZO-1, and Rho was analyzed by Western blot. GAPDH was used as a loading control. (F) Lung tissues were subjected to RT-PCR to examine to the mRNA levels of RhoA and RhoB. Data were expressed as the mean ± SD (n=3 in each group). ****P < .0001, ***P < .001, **P < .01 and *P < .05, Student’s t-test).

**Fig. 5.**
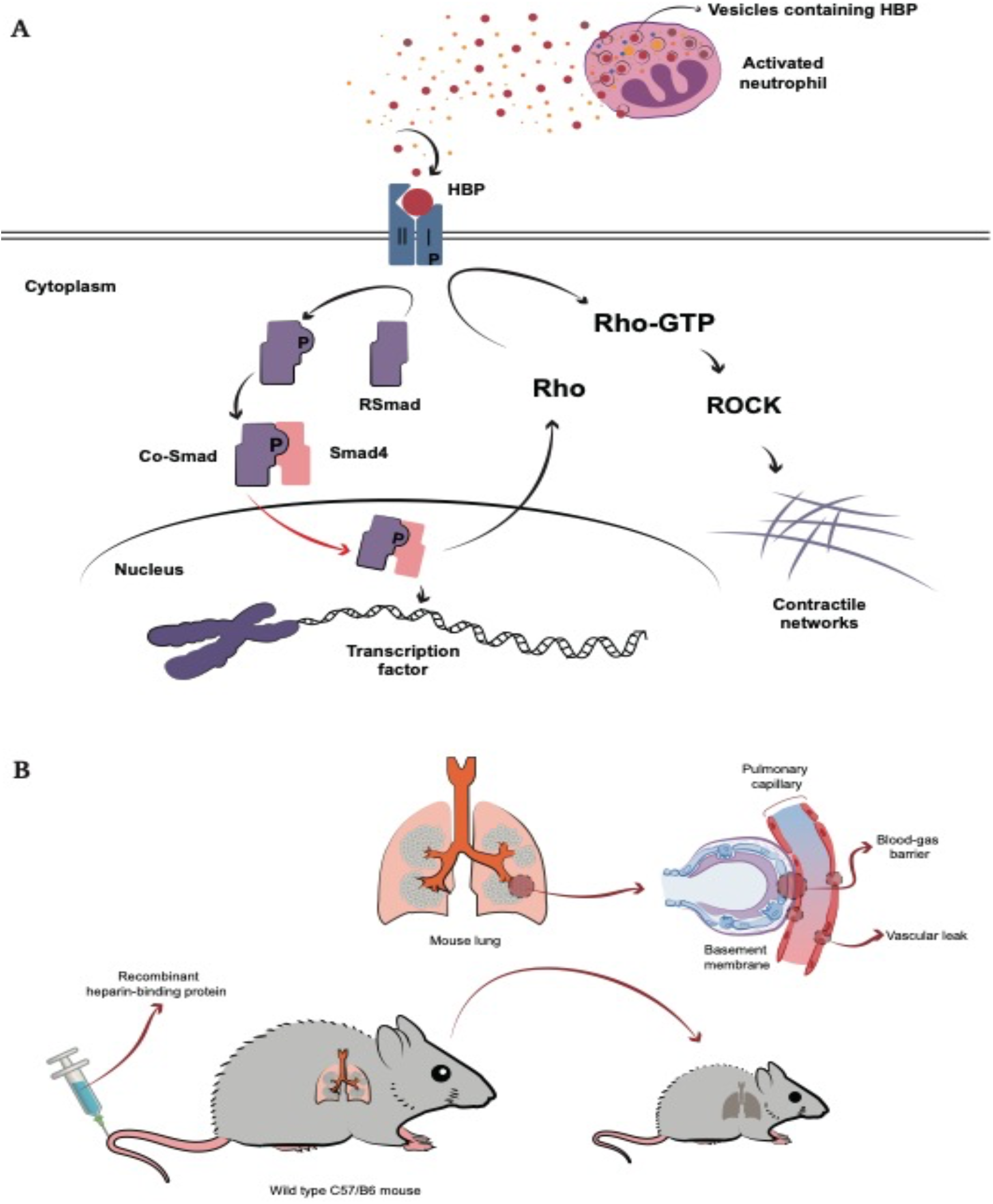
Proposed model of the role of HBP in TGF-β signaling pathway. The model is described in the text.

### HBP activated TGF-β/Rho signaling pathway in lung tissue in mice

Through the above experiments, we confirmed that HBP activated the TGF-β pathway by binding to TGF-β-R2 on the surface of HUVECs, resulting in cytoskeletal rearrangement and changes in tight junction proteins and thereby increased endothelial permeability. To further confirm the results *in vivo*, C57BL/6 mice were divided into four groups: control group, HBP group, TGF-β-R2 siRNA group, and TGF-β-R2 siRNA + HBP group. After the treatments, the animals were sacrificed, and the lung tissue was removed for related experiments. Western blot and Immunohistochemical analysis showed that the protein levels of phospho-SMAD2/3 in lung tissue were increased in HBP group, but total SMAD2/3 protein did not change significantly. TGF-β-R2 knockdown inhibited the effect caused by HBP treatment (Fig. 4E, Fig. S1A). We observed that the protein level of Rho was upregulated in lung tissue stimulated with HBP (Fig. 4E). Then we analyzed the effects of HBP treatment on the mRNA expression of RhoA and RhoB in lung tissue, the results showed that the mRNA levels of RhoA and RhoB were upregulated by HBP (Fig. 4F). Knockdown of TGF-β-R2 antagonized this effect.

## DISCUSSION

Through the above experiments, we reported that HBP as an extracellular ligand activated TGF-β signaling pathway by binding with TGF-β-R2 on the surface of ECs, thus increasing the permeability of ECs. To our knowledge, this is the first study to investigate the HBP-binding protein receptors on the endothelial surface. The effect of HBP on the blood–gas barrier *in vivo* was observed directly by TEM. We found that HBP treatment destroyed the integrity of tight junctions and induced the formation of intercellular gaps. Our study provides a new insight into the pathogenesis of HBP-induced ALI and the regulatory effect of HBP on endothelial cell permeability in sepsis.

Endothelial dysfunction plays a role in sepsis-induced organ dysfunction[22]. HBP was first discovered by Shafer et al. in 1984[23]. HBP has various functions, which can not only induce vascular leakage, edema formation, and leukocyte chemotaxis, but also promote the repairing process of tissues and cells by inhibiting caspase-3 activity[9] or inducing the migration of a series of cells [3, 10–13]. Although the mechanism of HBP-induced cell migration remains unclear, it is reasonable to assume that it may be related to the EMT process. It is widely known that EMT plays a fundamental role in many pathophysiological processes such as tissue repair, fibrosis, and tumor metastasis[24, 25], mainly through TGF-β signaling pathway. In addition, there is no direct experimental evidence for the existence of membrane receptors for HBP. Therefore, our study focused on the relationship between HBP and the TGF-βR/SMAD signaling pathway, confirming for the first time that HBP directly bound to the extracellular domain of TGF-β-R2 and activated TGF-β signaling pathway. However, we did not observe any significant changes in E-cadherin and N-cadherin upon HBP treatment (data not shown). One possible explanation is that HBP stimulation was not long enough to elicit EMT. It had been demonstrated that HBP can effectively inhibit apoptosis of HUVECs[9]. In our study, HUVECs were treated with HBP (0, 1, 10, and 50 μg/mL for 24 h), we found that HBP promotes HUVEC proliferation in a dose-dependent manner by CCK8 analysis (Fig. SB). We also observed that the mRNA levels of occludin and ZO-1 were slightly elevated upon HBP treatment (Fig. SC), which was consistent with the results conducted by Friedrich et al.[26]. Friedrich et al. showed that histone deacetylase inhibitors improved barrier recovery and the epithelial wound healing process via TGFβ1 signaling in inflammatory bowel diseases. We believe that HBP is a double-edged sword, which can not only cause inflammatory response and tissue damage due to its chemotactic properties, but also promote tissue repair through TGF-β signaling pathway.

We then attempted to determine the regulatory role of the TGF-βR/SMAD signaling pathway in HBP-induced injurious effects. At the cytoplasmic membrane level, TGF-βR-related ligands mediate the phosphorylation of downstream SMAD proteins and provide regulatory signals that control the expression of target genes. For example, TGF-β signaling activation has been found to upregulate RhoB expression and induce actin reorganization[18]. In addition to the SMAD-dependent pathway, TGF-β signaling activates Rho-ROCK pathway in a SMAD-independent way[27]. In this study, we observed that HBP-induced activation of TGF-β signaling pathway leads to a series of biological effects, including activation of the Rho-ROCK pathway, redistribution of tight junction proteins, and ultimately increased endothelial cell permeability. Furthermore, the viability of HUVECs increased upon HBP treatment via the TGF-β signaling pathway. Thus, we further elucidated the mechanism of HBP-induced vascular leakage by demonstrating that HBP acted as a ligand to TGF-β-R2 on the surface of ECs, thereby activating TGF-β signaling pathway.

Increased endothelial permeability leads to fluid leakage, which is involved in the pathogenesis of septic ALI/acute respiratory distress syndrome (ARDS). Under pathological conditions, fluid and proteins leak out of the blood and into the alveolar cavity owing to dysfunction of the blood–gas barrier. HBP has been reported to increase the permeability of ECs and is a potential mediator of sepsis-induced ALI[8]. Inhibition of HBP release can reduce pulmonary edema[28]. Moreover, HBP is considered as an important prognostic indicator of short-term mortality in septic ALI/ARDS[29]. To our knowledge, the changes in the blood–gas barrier and cell junctions in lung tissue upon HBP treatment have not been investigated. Using TEM, we observed that tight junctions between epithelial and endothelial cells opened and paracellular gaps were visible in HBP-treated lung tissue, indicating increased permeability of the blood–gas barrier. Notably, TGF-β-R2-siRNA antagonized the effects of HBP. Thus, our findings suggested that HBP induced ALI by activating TGF-β signaling pathway.

Previous study observed that normalization of plasma HBP level was associated with the recovery in septic patients. Therapeutic HBP may reduce vascular leakage and may be a potential therapeutic strategy for sepsis[30]. One way to decrease plasma HBP level is to block HBP release from neutrophils by using β2-integrin antagonist[31], simvastatin[30, 32], and tezosentan[33]. The other way is to block the effects of HBP on cells by using polyanion dextran sulphate[3], heparin, low molecular weight heparin[8], and bikunin[34]. In recent years, no specific receptor for HBP has been found. Fortunately, we identified the membrane receptor for HBP, which may direct new directions for treating sepsis.

The mechanisms of endothelial cell permeability are very complex. At present, many questions, such as the more specific mechanism of HBP-induced tight junction protein depolymerization and redistribution, remain unanswered. Whether HBP affects endothelial permeability through other signaling pathways remains to be further studied.

There are several limitations to this study. First, it is necessary to further analyze the causal link between ligand-receptor binding (HBP-TGF-β-R2) and the activation of downstream pathway. Second, the high cost of recombinant HBP limited the number and type of *in vivo* experiments that could be performed.

In summary, our study demonstrated that HBP bound to TGF-β-R2 as a ligand and activated TGF-β signaling pathway to induce ALI. HBP up-regulated Rho protein levels through the activation of TGF-β signaling pathway and activated the Rho-ROCK pathway to promote the cytoskeletal remodeling and the redistribution of tight junction protein (ZO-1), resulting in increased endothelial permeability.

## Materials and Methods

### DNA construction

RNA was extracted using Monzol (Monad, China) according to the manufacturer’s instructions. cDNA (RhoA, RhoB) was generated from 1 μg of each RNA sample (ShineGene, China). The sequences of the following primers were designed and purchased from Sangon Biotech (China), as shown in Table 1 (see supplemental information). Vector pGEX-4T-2 was used for the bacterial expression of proteins. The sequences used in human umbilical vein endothelial cells (HUVECs) and mice for expression of transforming growth factor-β receptor type 2 (TGF-β-R2) siRNA (small interfering RNA) were 5′-CCATCATCCTGGAAGATGA-3′ and 5′-GCTCGCTGAACACTACCAA-3′.

**Table 1.**
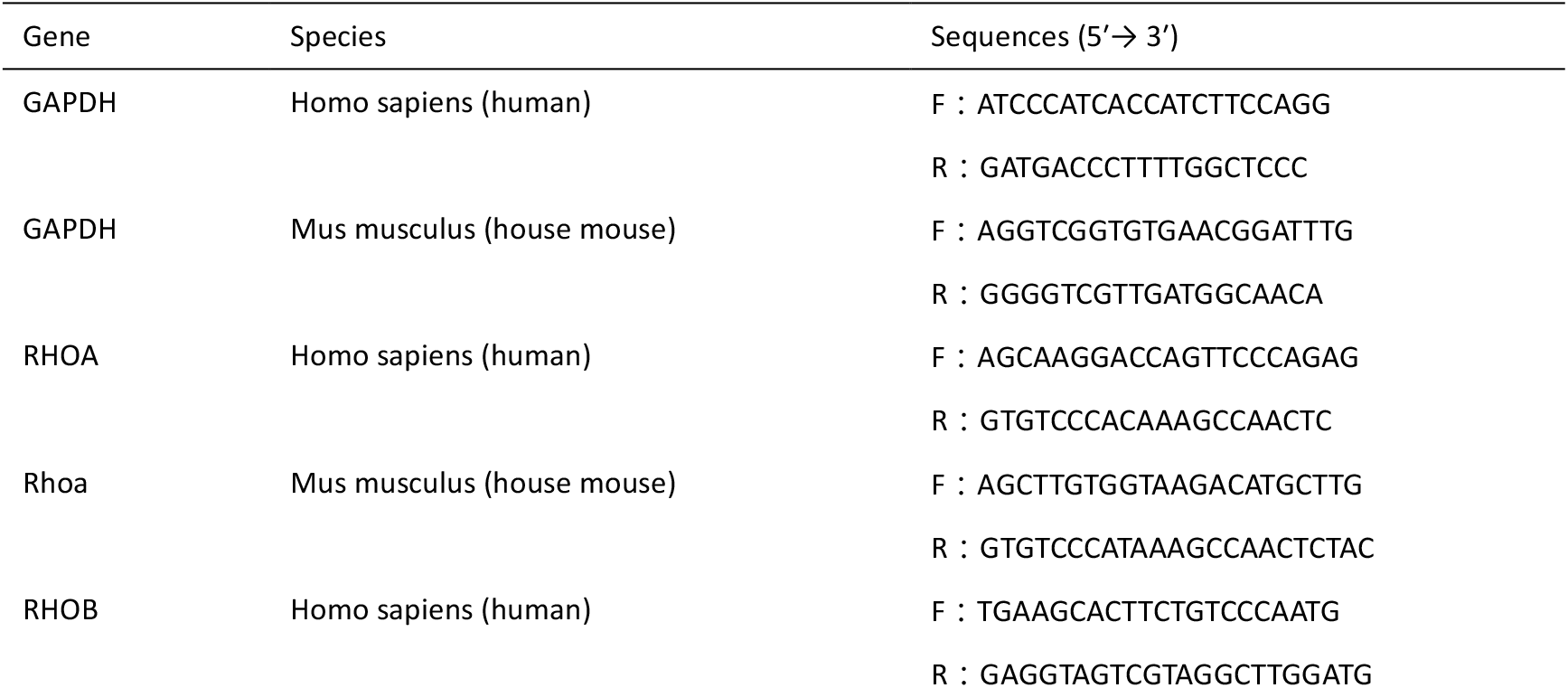
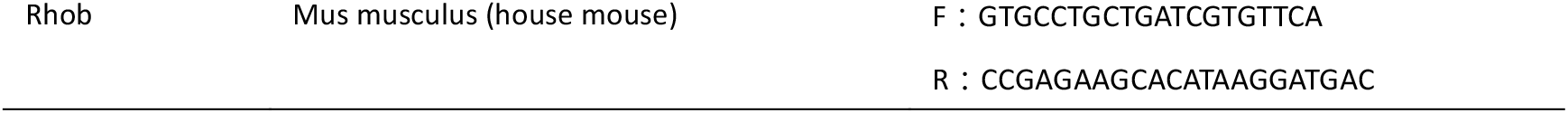
Primer sequences

### Antibodies and chemical reagents

In this study, we used anti-RHOA/RHOB/RHOC (mAb A4855, Abclonal), anti-ZO-1 (pAb A0659, Abclonal), anti-ZO-1 (D6L1E, mAb #13663, Cell Signaling Technology), anti-RHOB (pAb A2819, Abclonal), anti-phospho-SMAD2-S465/467+SMAD3-S423/425 (pAb AP0548, Abclonal), anti-RhoA (pAb A13947, Abclonal), anti-TGF-β-R2 (pAb A1415, Abclonal), anti-HBP (EPR9503, mAb ab181989, Abcam), anti-HBP (mAb #246322, R&D systems), anti-HBP (pAb ab167336, Abcam systems), anti-SMAD2/3 (D7G7, mAb #12470, Cell Signaling Technology), anti-SMAD2/3 (pAb AF6367, Affinity), goat anti-rabbit immunoglobulin G (IgG) H&L [horseradish peroxidase (HRP)] (ab27236, Abcam), anti-mouse IgG H&L (HRP) (ab27241, Abcam), Alexa Fluor 488 donkey anti-mouse (A21202, Thermo Fisher Scientific), Alexa Fluor 555 donkey anti-mouse (A31570, Thermo Fisher Scientific), Alexa Fluor 488 donkey anti-rabbit (A21206, Thermo Fisher Scientific), Alexa Fluor 555 donkey anti-rabbit (A31572, Thermo Fisher Scientific), 4′,6-diamidino-2-phenylindole; D1306, Thermo Fisher Scientific), Sera-MagTM SpeedBeads magnetic protein A/G particles (GE Life Sciences), and GST (glutathione S-transferase) Sepharose beads (17075601, GE Life Sciences). Recombinant human HBP (2200-SE-050, R&D Systems), protease inhibitor cocktail (HY-K0010), phosphatase inhibitor cocktail І (HY-K0021), phosphatase inhibitor cocktail ІI (HY-K0022), and Y27632 (HY-10071) were purchased from MedChem Express. Halofuginone (S8144, Selleck). Minute™ cytoplasmic and nuclear fractionation kit (SC-003, Invent Biotech). Cell counting kit-8 (M4839, AbMole).

### Cell culture and HBP treatment

Human umbilical vein endothelial cells (HUVECs) were purchased from Cyagen Biosciences. HUVECs were cultured in HUVECbasal medium (Cyagen Biosciences), supplemented with 10% (v/v) fetal bovine serum (FBS) (Cyagen Biosciences), 1% (v/v) penicillin-streptomycin (Cyagen Biosciences), 1% (v/v) glutamine, 1% (v/v) endothelial cell growth supplement (Cyagen Biosciences), and 1% (v/v) heparin (Cyagen Biosciences). Cells were cultured at 37°C in a humidified 5% CO2 incubator. Human recombinant HBP 10 μg/mL was used for *in vitro* experiments. Cells were washed three times with phosphate-buffered saline (PBS) and then treated with HBP for indicated time.

### SiRNA transient transfections

SiRNAs for human TGF-β-R2 (GenBank accession number NM_001024847.2, NM_003242.6) were synthesized by Guangzhou RiboBio (Guangzhou, China). SiRNA transient transfection was performed using the lipid-based method. For the lipofection procedure, jetPRIME (polyplus transfection, Illkirch, France) was used. A mixture of jetPRIME buffer (200 μL), siRNAs (50 nM), and JetPRIME reagent (4 μL) were directly added to HUVECs at 50% confluence. The medium was replaced with fresh medium 12 h after transfection. The efficiency of TGF-β-R2silencing was assessed by western blot analysis.

### Cell viability assay

A Cell Counting Kit-8 (CCK-8) assay (M4839, AbMole) was used to analyze cell viability. A total of 10^4^ cells/well were seeded into 96-well plates. The cells were divided into four groups. The cells were also stimulated with PBS or HBP (1, 10, 50 μg/mL) for 24 h and incubated with 100 μL of CCK-8 solution per well for 2 h. The cell viability was measured at 450 nm using a Tecan Microplate Analyzer (Spark 20M).

### Trans-endothelial permeability assays

HUVECs were grown on 0.4-mm pore Transwell filters (Fisher Scientific UK Ltd) until confluent. FITC-dextran was applied apically at 1 mg/mL and allowed to equilibrate for 30 min before a sample of the medium was removed from the lower chamber to measure permeability. Fluorescence was measured at 520 nm using a Tecan Microplate Analyzer (Spark 20M).

### Co-immunoprecipitation and immunoblotting assays

Cells were washed three times with PBS and lysed on ice with lysis buffer (P0013, Beyotime), containing 1 μmol phenylmethylsulfonyl fluoride and then centrifuged (4°C, 17,000 x g) for 15 min. Cell lysates were incubated with certain antibodies and 25 μL of magnetic A/G beads at 4°C for 12 h. Next, the protein-bead complexes were washed using lysis buffer and subjected to immunoblotting assay. The protein samples were electrophoresed on 8% or 10% precast sodium dodecyl sulfate–polyacrylamide gel and transferred onto polyvinylidene difluoride (PVDF) membranes (Millipore, USA). The PVDF membranes were incubated with Tris-buffered saline containing Tween (TBS-T) with 5% bovine serum albumin at room temperature for 1 h and then incubated overnight at 4°C with primary antibodies. After washing three times in TBS-T, the membranes were incubated for 60 min at room temperature with HRP-conjugated secondary antibody. The membrane was then washed in TBS-T for three times.

### GST pull-down assay

Bacterially expressed GST-TGF-β-R2, GST-TGF-β-R2 (23–166 amino acids), GST-TGF-β-R2 (167– 187 amino acids) and GST-TGF-β-R2 (188–567 amino acids) were purified using GST Sepharose beads in TNTE 0.5% buffer. His-HBP was purchased from R&D Systems. Purified His-HBP was incubated with beads bound with GST (control), GST-TGF-β-R2, GST-TGF-β-R2 (23–166 amino acids), GST-TGF-β-R2 (167–187 amino acids) and GST-TGF-β-R2 (188–567 amino acids) in TNTE 0.5% buffer at 4°C for 3 h. Finally, the protein-bead compounds were washed five times with the same buffer and subjected to western blot analysis.

### Immunofluorescence assay and confocal microscopy

Cells grown on glass coverslips were washed three times with PBS, fixed with 4% paraformaldehyde, and permeabilized with 0.3% Triton X-100. Next, the cells were stained with determined primary and proper fluorescently conjugated secondary antibodies. Alexa Fluor 555–conjugated phalloidin was used for F-actin staining. Images were obtained using a Nikon A1R confocal microscope using NIS-Elements AR software (Japan).

### Quantitative real-time polymerase chain reaction (qRT-PCR)

The samples were lysed and processed for total RNA extraction using Monzol according to the manufacturer’s instructions. The cDNA was synthesized by superscript reverse transcriptase. For normalization of the samples, the cDNA of the glyceraldehyde-3-phosphate dehydrogenase (GAPDH) gene was also amplified by the PCR instrument. The amplification reaction was carried out in a thermal cycler, and the value of 2(-⊿Ct) was calculated automatically by PCR instrument (FTC3000, Funglyn Biotech, Canada).

### Mice and HBP exposure protocol

Adult male C57BL/6 mice (6 weeks old, 25 ± 2 g,) were used. All the animals were maintained in the Experimental Animal Center of China Medical University, with a 12-h light/dark cycle with free access to food and water. The Council on Animal Care of China Medical University approved the experimental protocol. The mice were fasted overnight with free access to water before the experiment. The mice were randomly assigned to four groups (six per group): (1) control group: 10 mmol PBS (200 μL) intravenously; (2) HBP group: HBP 150μg/200 μL intravenously for 6 h; (3) TGF-β-R2 siRNA group: si-TGF-β-R2 20 μmol/100 μL intravenously for 48 h; (4) TGF-β-R2 siRNA + HBP group: 20 μmol/100 μL si-TGF-β-R2 intravenously for 48 h followed by HBP 150 μg/200 μL intravenously for 6 h. Anesthesia was initiated with 1.5% sodium pentobarbital (40 mg/kg, intraperitoneally). After the treatments, the animals were sacrificed by painless exsanguination after being anesthetized. Lungs were collected for further experiments.

### Lung wet/dry weight (W/D) ratio

The left lung lobes from random samples of four groups were resected at the end of the experiment and the wet weight was determined. The lungs were dried in an incubator at 60°C for 72 h before being weighed again to determine the dry weight. The W/D weight ratio was calculated as an indicator of lung edema.

### Histology and electron microscopy of lung

Lung tissues of the mice were soaked in 4% (v/v) paraformaldehyde for at least 24 h. After that, the tissues were dehydrated with graded alcohol, and then embedded in paraffin at 60°C. The specimens were cut into 5 μm sections and stained with hematoxylin and eosin (HE). The slide was examined using a light microscope (Nikon ECLIPSE 80i, Japan). Lung injury was scored according to ALI scoring system[35]. Tissue samples for electron microscopy were fixed in 2.5% glutaraldehyde at 4°C overnight and washed with PBS (0.1 M). Next, the tissue samples were fixed with 1% osmic acid for 1 h. The tissues were dehydrated with graded alcohol and acetone. The tissue samples were then permeated with Epon812 and embedded at 60°C for 48 h. The samples were sliced into 70-nm sections and stained with uranyl acetate and lead citrate. The specimens were examined with a transmission electron microscope (TEM) (HITACHI H-7650) operated at an acceleration voltage of 15 kV at a ×20,000–60,000 magnification. The specimens were also examined with a scanning electron microscope (SEM) (HITACHI SU3500) operated at an acceleration voltage of 15 kV, working distance of 5 mm, and ×1000–5000 magnification.

### Statistical analysis

All data were processed using SPSS version 22.0 software and GraphPad Prism 8 software. Data were expressed as mean ± standard deviation (SD). Student’s t-test was used to determine the significant difference between the two groups. A P value of < 0.05 was considered statistically significant.

## Abbreviations

HBP: Heparin-binding protein
TGF-β-R2: Transforming growth factor-β receptor type 2
EMT: Epithelial–mesenchymal transition
HUVEC: Human umbilical endothelial cell
PMN: Polymorphonuclear neutrophil
SEM: Scanning Electron Microscope
TEM: Transmission Electron Microscope
ZO: zonula occludens
GAPDH: glyceraldehyde-3-phosphate dehydrogenase
GST: Glutathione S-transferase.

## ACKNOWLEDGEMENTS

This study was supported by grants from the clinical research center of critical care medicine, Shenyang (Grant NO. 20-204-4-41) and clinical research center of critical care medicine, Liaoning province. Thanks to Professor Liu Cao of Department of Life Science, China Medical University, for his guidance on this experiment.

## SUPPLEMENTARY INFORMATION

### Availability of data and materials

The data used to support the findings of this study are included within the article.

### Funding information

Clinical research center of critical care medicine, Shenyang, Grant Number: 20-204-4-41. Clinical research center of critical care medicine, Liaoning province.

## Declarations

### Ethics approval and consent to participate

The animal study was reviewed and approved by China Medical University Institutional Animal Care and Use Committee.

### Consent for publication

Not applicable.

---

**Fig. S.**
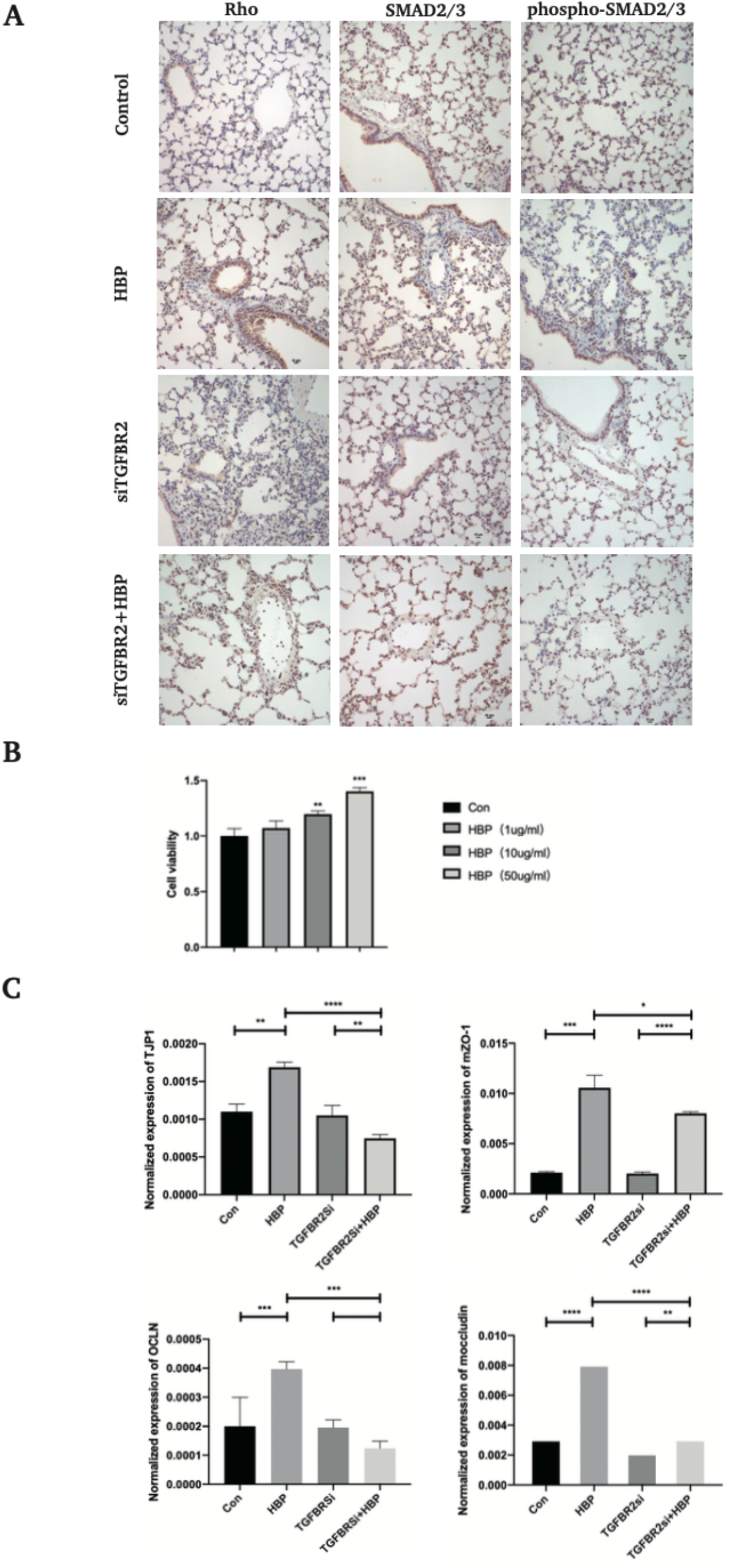
(A) TGF-β-R2-siRNA transfected C57BL/6 mice were treated with HBP intravenously at the concentration of 150 μg/mouse for 6 h. After the treatments, the lung tissue was taken for immunochemistry assay. Protein expression and distribution of Smad2/3, phospho-Smad2/3, and Rho in the lung tissues were observed by immunochemistry assay (original magnification × 400). (B) HUVECs were treated with HBP at the concentrations of 0, 1, 10 and 50 μg/mL for 24 h, and cell viability was determined by CCK8 assay. TGF-β-R2-siRNA HUVECs were treated with HBP for 12 h at the concentration of 10 μg/mL. (C) HUVECs and mouse lung tissue were then subjected to qPCR assay to examine ZO-1 and occludin. Data were expressed as the mean ± SD (****P < .0001, ***P < .001, **P < .01 and *P < .05, Student’s t-test).

